# Transcutaneous vagus nerve stimulation in humans induces pupil dilation and attenuates alpha oscillations

**DOI:** 10.1101/2020.08.25.265876

**Authors:** Omer Sharon, Firas Fahoum, Yuval Nir

## Abstract

Vagus nerve stimulation (VNS) is widely used to treat drug-resistant epilepsy and depression. While the precise mechanisms mediating its long-term therapeutic effects are not fully resolved, they likely involve locus coeruleus (LC) stimulation via the nucleus of the solitary tract (NTS), which receives afferent vagal inputs. In rats, VNS elevates LC firing and forebrain noradrenaline levels, whereas LC lesions suppress VNS therapeutic efficacy. Non-invasive transcutaneous VNS (tVNS) employs electrical stimulation that targets the auricular branch of the vagus nerve at the cymba conchae of the ear. However, the extent that tVNS mimics VNS remains unclear. Here, we investigated the short-term effects of tVNS in healthy human male volunteers (n=24), using high-density EEG and pupillometry during visual fixation at rest. We compared short (3.4s) trials of tVNS to sham electrical stimulation at the earlobe (far from the vagus nerve branch) to control for somatosensory stimulation. Although tVNS and sham stimulation did not differ in subjective intensity ratings, tVNS led to robust pupil dilation (peaking 4-5s after trial onset) that was significantly higher than following sham stimulation. We further quantified, using parallel factor analysis, how tVNS modulates idle occipital alpha (8-13Hz) activity identified in each participant. We found greater attenuation of alpha oscillations by tVNS than by sham stimulation. This demonstrates that tVNS reliably induces pupillary and EEG markers of arousal beyond the effects of somatosensory stimulation, thus supporting the hypothesis that tVNS elevates noradrenaline and other arousal-promoting neuromodulatory signaling, and mimics invasive VNS.

**Significance statement:** Current non-invasive brain stimulation techniques are mostly confined to modulating cortical activity, as is typical with transcranial magnetic or transcranial direct/alternating-current electrical stimulation. Transcutaneous vagus nerve stimulation (tVNS) has been proposed to stimulate subcortical arousal-promoting nuclei, though previous studies yielded inconsistent results. Here we show that short (3.4s) tVNS pulses in naïve healthy male volunteers induced transient pupil dilation and attenuation of occipital alpha oscillations. These markers of brain arousal are in line with the established effects of invasive VNS on locus coeruleus-noradrenaline signaling, and support the notion that tVNS mimics VNS. Therefore, tVNS can be used as a tool for studying the means by which endogenous subcortical neuromodulatory signaling affects human cognition, including perception, attention, memory, and decision-making; and also for developing novel clinical applications.

## Introduction

Since 1988, vagus nerve stimulation (VNS) has been successfully used to reduce epileptic seizures in patients with drug-resistant epilepsy (Krahl and Clark, 2012), and has demonstrated clinical effectiveness for many patients treated with invasive VNS (Boon et al., 2018; Kwon et al., 2018). VNS is also applied as a treatment for drug resistant major depression (e.g Nemeroff et al., 2006).

VNS modulates vagal afferent inputs to the brainstem Nucleus Tractus Solitaris, which subsequently activate the locus coeruleus-noradrenaline (LC-NE) system. Indeed, in rats, VNS increases LC neuronal discharges (Takigawa and Mogenson, 1977; Groves et al., 2005; Hulsey et al., 2017) and elevates NE levels in the hippocampus and cortex (Dorr and Debonnel, 2006; Roosevelt et al., 2006). The effects of VNS on LC-NE are considered key to reducing seizures. This is due to the strong positive correlation observed of the noradrenergic and anticonvulsive effects of VNS (Raedt et al., 2011), and due to the elimination of the anticonvulsive effects by means of the chemical lesions of the LC (Krahl et al., 1998). VNS also modulates signaling in other neuromodulatory pathways such as the serotonergic, dopaminergic, and cholinergic systems (Dorr and Debonnel, 2006; Manta et al., 2009; Mridha et al., 2019). However, some of these effects are likely to be secondary, i.e. occur later and with mediation through the LC-NE system (Dorr and Debonnel, 2006).

In humans, invasive VNS induces markers of brain arousal that are consistent with LC-NE activity. This includes pupil dilation (Desbeaumes Jodoin et al., 2015), which is tightly linked with LC-NE activity (Joshi et al., 2016; Reimer et al., 2016; Gelbard-Sagiv et al., 2018; Hayat et al., 2020). VNS may also lead to EEG desynchronization, but the effects are subtler than in pupil dilation, at least regarding the clinical parameters that typically employ long (30-60s) stimulation epochs. Accordingly, early studies with <10 patients each, did not find VNS effects on spontaneous intracranial EEG (Hammond et al., 1992) or scalp EEG (Salinsky and Burchiel, 1993). In contrast, a more recent study with 19 participants, which analyzed separately VNS ‘responders’ and ‘non-responders’, observed EEG desynchronization in the alpha and delta bands (Bodin et al., 2015).

Non-invasive transcutaneous vagal nerve stimulation (tVNS) applies electrical current at a high frequency (typically 25Hz) through the left ear, targeting the auricular branch of the vagus nerve at the cymba conchae (Figure 1) (for anatomic evidence see Van Bockstaele et al., 1999; Bermejo et al., 2017). tVNS has been shown to mimic the anticonvulsive and antidepressant effects of invasive VNS (Stefan et al., 2012; He et al., 2013; Hein et al., 2013; Bauer et al., 2016; Rong et al., 2016; Trevizol et al., 2016), and has demonstrated safety and tolerability (Redgrave et al., 2018). Beyond the clinical efficacy of tVNS, interest has grown regarding its use in healthy individuals for basic neuroscience research (Van Leusden et al., 2015). However, the literature is inconsistent as to the extent that tVNS mimics the effects of invasive VNS on EEG or pupil dilation; such evidence would suggest LC-NE involvement (Ventura-Bort et al., 2018; Warren et al., 2018; Keute et al., 2019). We suspect that the discrepancies stem from employing long (e.g. 30sec) stimulation epochs as in clinical applications, and due to the indirect focus on the P300 component in which LC-NE activity is assumed to play a key role.

**Figure 1:**
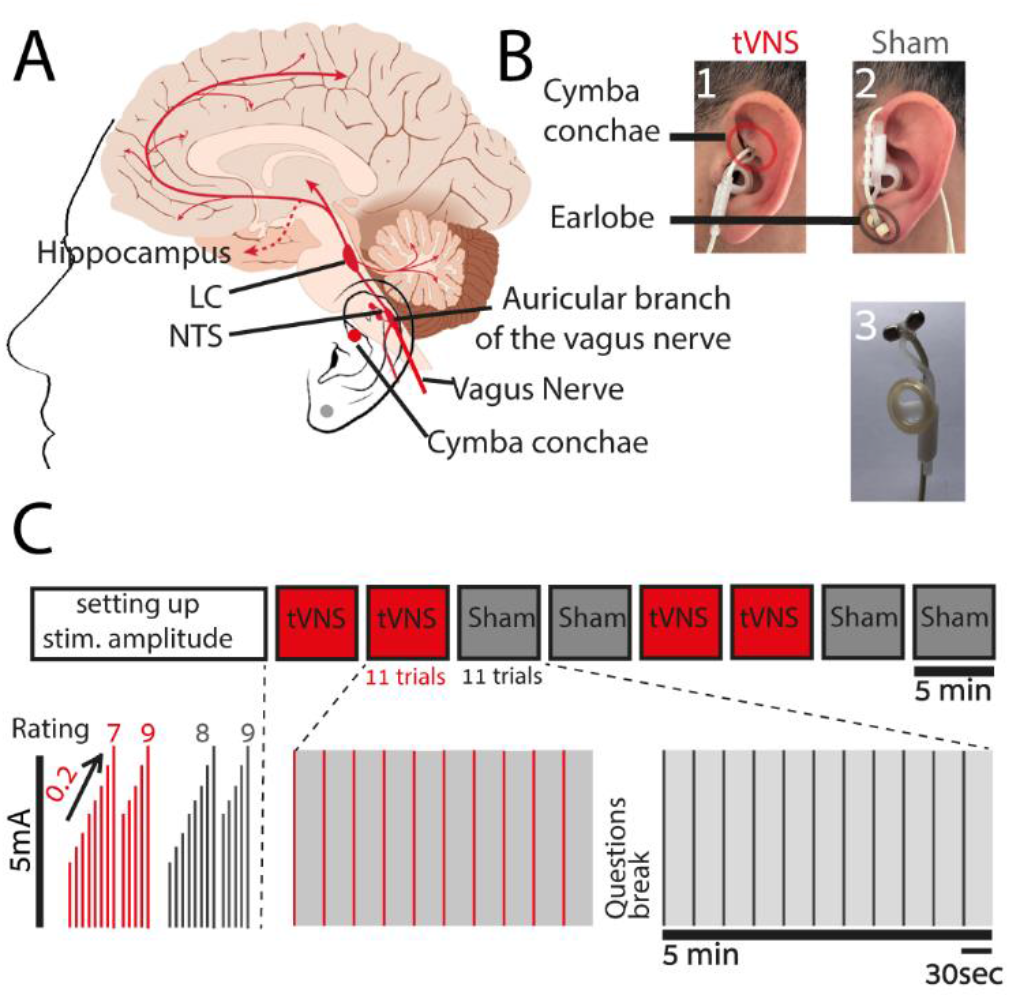
Experimental design. **Legend**: (A) Schematic illustration of the rationale of tVNS (B) Stimulation electrode placement – (1) location of the tVNS on the cymba conchae of the left ear (2) location of the sham stimulation on the left earlobe (3) photo of the commercial stimulation electrode. (C) Experimental design, each experiment started with a ‘method of limits’ procedure in order to adjust the stimulation current according to the individual subjective pain report (Rating); and then increased incrementally by 0.2 mA until a current matched to a rating of 8 was selected. Eight blocks were then conducted, each of 5min and including 11 stimulation trials of 3.4s and stimulation intervals of 25-27s.

Here, we set out to examine if short-term tVNS induces EEG and pupillary markers of arousal, as is in the context of VNS-induced activation. We used short (3.4s) stimulation pulses during task-free rest conditions in healthy naïve male volunteers (to avoid long-term changes associated with therapeutic effects (Follesa et al., 2007; Manta et al., 2013)). We hypothesized that if indeed tVNS increases LC and neuromodulatory activities, it should lead to pupil dilation, as has been observed across multiple species (Joshi et al., 2016; Reimer et al., 2016; Hayat et al., 2020). In addition, we hypothesized that tVNS would attenuate alpha oscillations that are anti-correlated with arousal during rest (Torsvall and Akerstedt, 1987a; Drapeau and Carrier, 2004a; Amzica and Lopes da Silva, 2017), and that are attenuated by invasive VNS (Bodin et al., 2015). In line with these predictions, we found that tVNS induces pupil dilation and alpha desynchronization above and beyond the effects of sham (somatosensory) stimulation.

## Materials and methods

### Participants

High-density (256-channel) EEG and pupillometry were recorded in 25 healthy young male adults (mean age: 28.08 ±5.84 years, 2 left-handed). Written informed consent was obtained from each participant. The study was approved by the Medical Institutional Review Board (IRB) at the Tel Aviv Sourasky Medical Center. Females of child bearing age were not included, per guidelines of the approved IRB. Participants reported being healthy and without a history of neuropsychiatric disorders; they indicated their dominant eye for pupillometry. One participant was excluded from the analysis due to excessive blinking, after which 24 participants remained (mean age: 28.3 ± 1.2). Data from an additional three participants were excluded from the EEG analysis due to lack of alpha activity, after which 21 participants remained for the EEG analysis (mean age 28.01±1.3).

### Experimental design

#### Main experiment

After the EEG setup (see below), participants performed a short ‘method of limits’ procedure to select tVNS/sham stimulation intensities while sitting. This procedure systematically identifies the maximal comfortable stimulation levels for each individual, as in (Kraus et al., 2013; Yakunina et al., 2017; Ventura-Bort et al., 2018). We applied 5s-long stimulation trials, starting at 0.1mA, and increasing in each trial by 0.2mA. After each trial, participants rated the subjective intensity on a scale of 0-9 ([0] = no sensation; [3] = light tingling; [6] = strong tingling; [9] = painful). We continued increasing the current until either reaching a level rated as [9] or a maximal level of 5mA. This procedure was carried out twice for each stimulation location (real tVNS at the cymba conchae vs. sham stimulation at the ear lobe). The mean currents corresponding to a subjective rating of [8] (just below painful) were selected, for each stimulation location separately. Thus, tVNS intensity was adjusted for each participant and location separately, as above the detection threshold and below the pain threshold, as is done in clinical settings (Ellrich, 2011). Participants were then instructed to position their heads in a chin-rest apparatus for adjusting and calibrating the eye-tracker (see below). Subsequently, participants were instructed to fixate on a white cross on a background of a gray computer screen (HP model 2311x, positioned 80cm from the participants’ eyes), throughout experimental “blocks” lasting 5 minutes. Each block included 11 trials of 3.4s stimulation epochs (in each trial, tVNS intensity was ramped up gradually, to the level defined above), separated by inter-stimulus-intervals of 26s (±1s jitter). We performed two blocks of either tVNS or sham, and then switched to position the stimulating electrode in the alternate location (the order was counterbalanced), to reach a total of eight blocks per session (Figure 1C). Before changing the electrode location, participants answered questions regarding their subjective experience of stimulation (Table 1). Participants were free to rest between the blocks ad lib. Data acquired during these “breaks” were used to characterize alpha activity in each individual during non-experimental conditions (see below).

**Table 1.** Subjective ratings of tVNS/sham stimulation.

|  | tVNS | sham |
| --- | --- | --- |
| I have a headache | $1.82 \pm 0.20$ | $1.89 \pm 0.22$ |
| I feel nausea | $1.67 \pm 0.19$ | $1.63 \pm 0.19$ |
| I feel dizziness | $1.82 \pm 0.22$ | $1.97 \pm 0.25$ |
| I feel neck pain | $1.87 \pm 0.16$ | $1.97 \pm 0.19$ |
| I feel muscle contraction in the neck | $2.10 \pm 0.20$ | $2.00 \pm 0.21$ |
| I feel stinging under the electrode | $2.22 \pm 0.22$ | $2.18 \pm 0.23$ |
| I feel skin irritation in the ear | $2.17 \pm 0.28$ | $2.15 \pm 0.27$ |
| I feel fluctuation in concentration | $4.02 \pm 0.39$ | $3.73 \pm 0.36$ |
| I have an unpleasant feeling | $2.52 \pm 0.29$ | $2.71 \pm 0.31$ |
| I am in a good mood | $4.82 \pm 0.32$ | $5.23 \pm 0.27$ |
| I am alert | $4.07 \pm 0.32$ | $4.15 \pm 0.31$ |

#### Pilot experiment

A similar experiment using the same device with a separate group of 29 male participants (mean age: 26.82 ±1.1 years, 4 left-handed) used the default clinical stimulation mode (30s on, 30s off) during fixation at rest, while recording high-density EEG (n=15) and pupillometry (n=29).

### Transcutaneous vagus nerve stimulation (tVNS)

tVNS was delivered using NEMOS®, (Cerbomed, Germany, now tVNS technologies; Figure 1B). In the tVNS condition, the electrodes were placed at the left cymba conchae, which is heavily innervated by the auricular branch of the vagus nerve (Peuker and Filler, 2002; Safi et al., 2016; Badran et al., 2018) (Figure 1A). In the sham condition, the electrodes were placed at the left earlobe (Figure 1A), which is not expected to induce brainstem or cortical activation (Kraus et al., 2007; Sellaro et al., 2015; Steenbergen et al., 2015). Pulses (200-300μs width) were delivered at a rate of 25Hz (duty cycle of ~7% ON time) for 3.4s. This included ramping up of intensity (as set by the device) to a level experienced as just-below painful, adjusted for each participant and condition separately (‘method of limits’ procedure above), as is often set clinically in patients (Vonck et al., 2014). To achieve 3.4s stimulation trials, we controlled the NEMOS stimulation device using linear actuators (Actounix, Canada) that pressed the ON/OFF button automatically according to programmable times. These actuators were controlled by Arduino mega (Arduino, Italy), directed by Psychopy python package (Peirce, 2007). Two additional measures verified good electrode contact throughout, and consistent effectiveness of the stimulation: (i) the NEMOS device stops stimulation automatically whenever good physical contact with the participant’s skin is disrupted, and (ii) the experimenter verified in each participant the presence of a visually-evident 25Hz stimulation artifact in EEG electrodes close to the left ear.

### Pupillometry

#### Data acquisition

Eye movements/gaze and pupil size were recorded monocularly from the dominant eye using an infrared video-oculographic system with a chin-rest (Eyelink 1000 Plus, SR Research). Gaze and pupil data were sampled at 500Hz, and positions were converted to degrees of visual angle based on a 9-point calibration performed at the beginning of the experiment (on mid-gray background). The experiment was carried out in a room with constant ambient light.

#### Data analysis

Pupil data were low-pass filtered, using a 10Hz 4th-order Butterworth filter with a zero-phase shift. Periods of blinks were detected using the manufacturer’s standard algorithms, with default settings. The remaining data analyses were performed using custom-made Matlab scripts (The MathWorks). Blinks were removed by linear interpolation of values measured 100ms before and after each identified blink (de Gee et al., 2014). Peri-trial data were segmented by extracting pupil data [−10s +13.4s] around each stimulation trial. Trials in which interpolated data accounted for > 50% of the data points were excluded (van Steenbergen and Band, 2013). After excluding one participant who had no trials remaining, the process yielded a mean of 42.12±1.79 trials in the tVNS condition and 42.16±1.79 in the sham condition (of 45, range 42-45 for both). To enable averaging across participants with different pupil sizes while avoiding arbitrary units, we converted pupil data to ‘percent change’ values relative to a 10s baseline prior to stimulation: [(x−baseline/baseline)*100], as in (Reimer et al., 2016; Liu et al., 2017). Baseline pupil values did not differ significantly between the tVNS and sham conditions. In both conditions, the smaller the pupil before a specific trial, the higher the chance of observing significant pupil dilation (R=−0.27, p<10-20, see Discussion).

The resulting pupil time-courses were the mean values across trials for each participant and condition separately, as depicted in Figure 2A (for visualization only, single participant traces were band-pass filtered again between 0.01-10Hz, as shown in Figure 2C). To present the individual participant data, we reduced the pupil data for each participant and condition to a scalar value (Figure 2B), by averaging the time-course across trials in the interval between the two points of half maximum (FDHM, 3.2-10.4s) following stimulation onset (see the dashed bar in Figure 2A).

**Figure 2:**
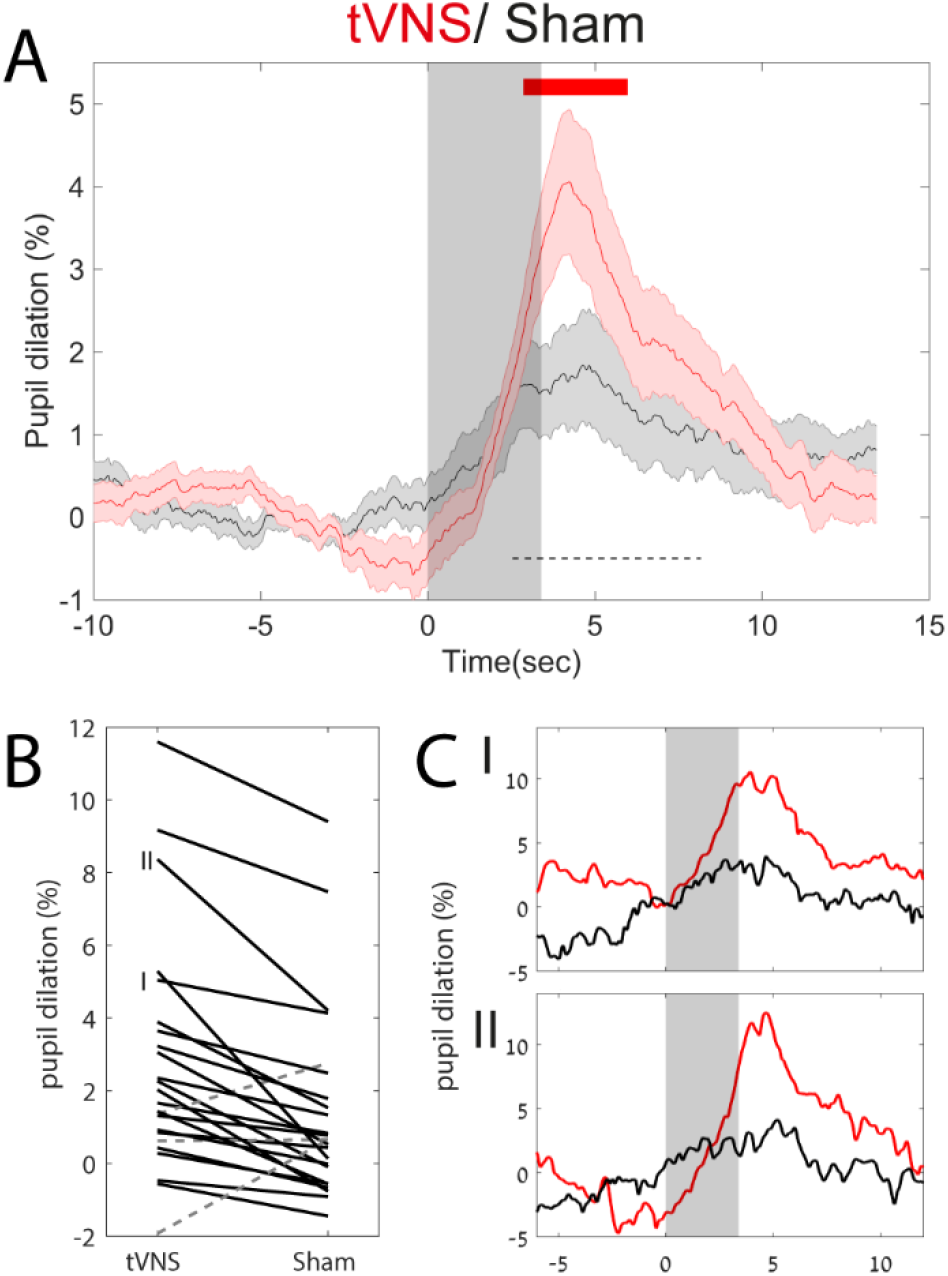
tVNS leads to greater pupil dilation than sham stimulation. **Legend**: **(A)** Grand average pupil dilation in response to tVNS (red trace) and sham stimulation (black trace). Shaded areas around the trace indicate the standard error of the mean. The grey transparent rectangle indicates that the active current is on. The upper red line indicates FDR-corrected statistical significance using the Wilcoxon signed rank test. The dashed black bar indicates the time interval used to compute individual subject dilation values in B. **(B)** Single participant values in both tVNS and sham conditions between the 2 points of half maximum (FDHM, 3.2-10.4s dashed black bar in A). The solid black lines denote tVNS>sham, while the dashed grey lines denote sham>stVNS. I,II refers to the single participant traces shown in C. **(C)** Two representative single-subject pupil time-courses as indicated in B, with identical graphic representation as in A.

Gaze data and blink rate were also inspected and compared between conditions. Gaze was extracted, interpolated, and averaged using the same procedure described above for pupil size. Data points marked as blinks were summed across participants to produce a blink rate that was time-locked to stimulation onset.

### EEG

#### Data acquisition

High-density EEG was recorded continuously using a 256-channel hydrocel geodesic sensor net (Electrical Geodesics, Inc. [EGI], Eugene OR, USA). Each carbon-fiber electrode, consisting of a silver chloride carbon fiber pellet, a lead wire, and a gold-plated pin, was injected with conductive gel (Electro-Cap International). Signals were referenced to Cz, amplified via an AC-coupled high-input impedance amplifier with an antialiasing analog filter (NetAmps 300, EGI), and digitized at 1000 Hz. Electrode impedance in all sensors was verified to be <50 kΩ before starting the recording.

#### EEG data analysis

EEG preprocessing was performed in Matlab using custom-written code and the FieldTrip toolbox (Oostenveld et al., 2011). First, we used a subsample of 192 electrodes placed directly on the skull (avoiding cheek electrodes with higher muscle artifacts). Continuous data from these electrodes were segmented to 33s epochs, [−15s +18s] around each stimulation onset. To enable effective visual inspection, data epochs were initially de-trended linearly, notch-filtered (at 50Hz), and high-pass (>0.1Hz) filtered using a 2nd order Butterworth filter to remove DC shifts. We then visually confirmed that all sham and tVNS trials showed 25Hz stimulation artifact around the left ear. Trials without the artifact were excluded (a mean of 14.75%±3.08 trials were excluded). To focus on alpha oscillations, data were further band-passed filtered at two frequencies. The first filter was applied at 5-15Hz using a 3rd-order two-pass Butterworth filter, as in previous parallel factor analysis (PARAFAC) studies (Barzegaran et al., 2017). An additional notch filter at 25Hz (stimulation frequency) was used, with harmonics up to 475Hz, to remove any residual artifact stemming from stimulation and not removed by previous filters. Then, we removed the minimal number of channels or trials whose data crossed an absolute amplitude threshold of 100μV in an automatic iterative process – that is, each 3s epoch in each channel had a Boolean value [max(abs(x))>100]. Subsequently, in each iteration, either a channel or a trial was excluded, such that a minimal number of channel×trial 3s data epochs was discarded (the code is available at: https://github.com/sharomer/eeg_2d_minimal_rejection). This process removed large movement artifacts, but not all blink artifacts, which were separated later using the parallel factor analysis.

This preprocessing resulted in identifying a mean number of 18.76±2.86% unacceptable channels per participant (of 192, data were interpolated using a linear distance weighted interpolation), and a mean number of 22±3.42% unacceptable trials per participant (discarded from subsequent analyses). Only then, were trials classified to the tVNS or sham condition, to avoid any bias in preprocessing. The mean number of valid trials in the tVNS condition was 35.61±1.09, and in the sham condition, 35.61±1.09 (of 44, the number of trials did not differ significantly between the conditions). Next, data of each trial were transformed to the time-frequency domain using the Fast Fourier Transform (FFT), after multiplying by a moving hamming window of 3s. This yielded a frequency resolution of 0.33Hz and a temporal resolution of 0.33s.

#### Parallel factor analysis

We first extracted data from “break” periods (between stimulation blocks) to identify each participant’s alpha topography and frequency in an unbiased manner with respect to the study objectives. These data were segmented to 5s epochs, with 1s overlap with the preceding epochs and 1s overlap with the subsequent epochs. The epochs were band-passed filtered (as described for stimulation data, above) and reduced to 3s trials (discarding the overlap) to avoid filtering artifacts at the edges. Then, “break” data epochs were cleaned as described for stimulation data, using the same procedure described above (resulting in 117.84±7.15 trials, on average, per participant, with 175.28±2.71 clean channels on average). These 3s time-frequency epochs were used to identify each participant’s alpha topography and precise frequency range using the PARAFAC analysis (Harshman, 1970), as implemented in the N-way toolbox (Andersson and Bro, 2000), and as presented in Figure 3. The type of constraint for each dimension was set to non-negativity. The proper number of components was determined by using the Core Consistency Diagnostic (CCD), in which the number of components is the highest when the minimal value of CCD is 55% and 90.60±3.18% on average (Bro and Kiers, 2003).

**Figure 3.**
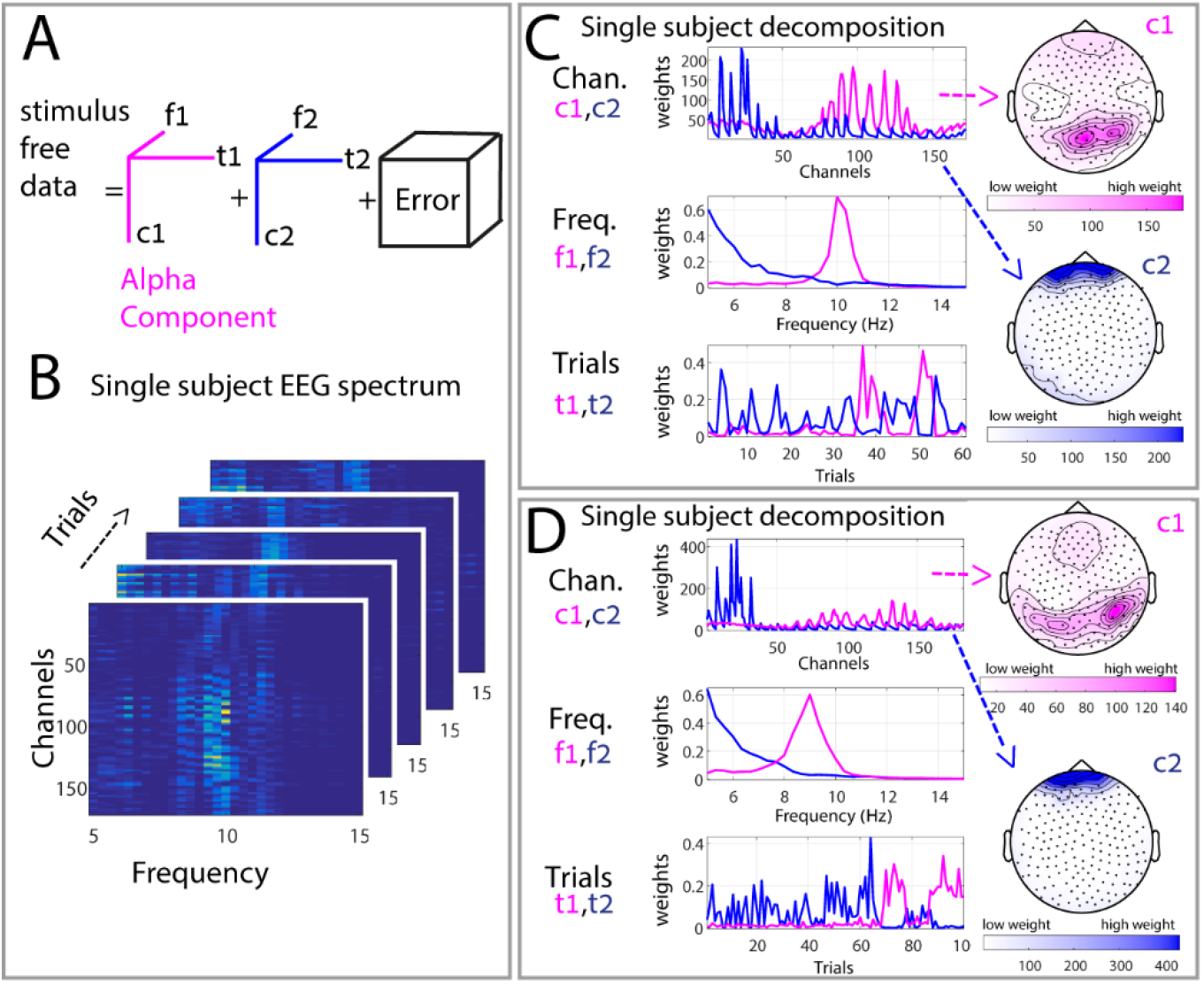
Parallel Factor Analysis (PARAFAC) to identify individual alpha activity. **Legend**: Graphic illustration of the Parallel Factor Analysis (PARAFAC) method we used to decompose the stimulus-free (break) data and create subject-specific topographical and frequency bands of interest. (A) Illustration of the PARAFAC model with two components, in which f1 and f2 refer to the frequency features, t1 and t2 indicate temporal features, and c1 and c2 represent the spatial features of the components in the channel space. (B) Spectrogram of five single 3s “trials” derived from the break, the same subject as in the top left in C. (C,D) Representative examples of the decomposition result for two participants. Each panel includes two components:1 (pink), and 2 (blue), together with their associated frequency (f) and trial (t) profiles. The spatial (channel) dimension is presented as scalp topographies on the right side.

Next, to assess the changes in alpha oscillations during stimulation, the individual weights for alpha component topography and frequency (Figure 4A) were derived from the break data, and multiplied by the spectrum of all channels, such that a single channel representing the weighted activity was achieved. We then subtracted the mean baseline activity in [−1:0]s relative to stimulation onset, for each participant and for each trial, and calculated the mean activity across participants (N=21); this yielded the results depicted in Figure 4D.

**Figure 4.**
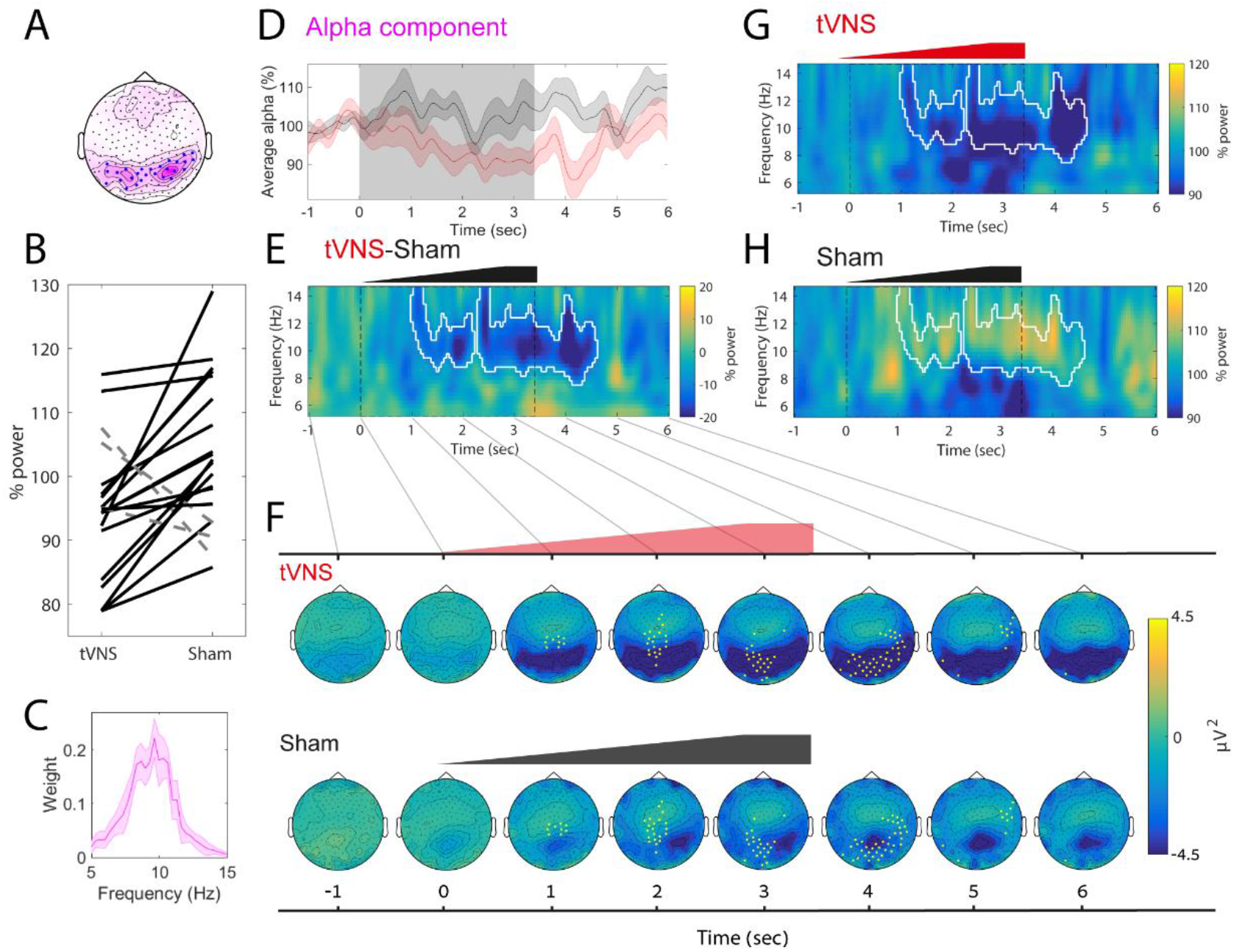
tVNS leads to greater attenuation of EEG alpha activity than does the sham stimulation. Legend: (A) Median alpha component topography. The median weights across participants are colored pink. The blue points mark electrodes with the highest alpha activity (selected using a threshold applied to the median weights) to facilitate visualization in subsequent panel E, but these electrodes are not used in any statistical analyses. (B) Alpha attenuation relative to baseline in individual subject data between 0:4sec, using the weighted topography in A and using the spectral profile in C. Black solid lines mark participants with higher alpha decreases in the tVNS condition (19/21), whereas dashed gray lines mark participants with higher alpha decreases in the sham condition (2/21). (C) Alpha component spectral profile (median across participants). (D) The mean alpha component time-course (using the spectral profile depicted in C, and the topographical profile depicted in A). (E) The difference in induced power between the tVNS and sham conditions (shown separately in G and H). White contours mark statistically significant time-frequency clusters (after correction for multiple comparisons). Note that tVNS causes alpha attenuation lasting several seconds. (F) Topographical dynamics following stimulation (at a resolution of 1s) reveals occipital alpha attenuation upon tVNS (upper panel) but not in the sham condition (lower panel). The yellow points mark electrodes comprising the statistically significant time-space cluster that exhibits tVNS attenuation > sham attenuation (after correction for multiple comparisons). (G) Mean induced spectrogram upon tVNS; the white contour is identical to that shown in E. (H) The mean induced spectrogram upon sham stimulation; the white contour is identical to that shown in E.

To assess more carefully the brain activity following stimulation, beyond the a-priori electrode- and frequency-band of interest, we conducted the following analyses. (i) We rigidly set the alpha topography (to investigate time-frequency changes in the entire spectrogram). To this end, we used the topography of interest derived from the PARAFAC decomposition of the break data (Figure 4A, lower panel), ignored the frequency of interest, plotted the entire spectrogram at 5-15Hz in % change relative to the same baseline ([−1:0]s prior to stimulation onset, Figure 4G,4H,4E), and used a cluster permutation test (see below). We also confirmed differences between tVNS and sham conditions by means of a post-hoc direct comparison using Wilcoxon signed rank tests (Results). (ii) Alternatively, we rigidly set the frequency-band of interest (to investigate changes in all electrodes). To this end, we used the frequency-of-interest derived from the PARAFAC decomposition of the break data (Figure 4A) and ignored the topography of interest. We plotted the entire topographical changes in voltage around times of stimulation, while subtracting the activity [−1:0s] prior to stimulation (Figure 4F). We then performed a topographical cluster permutation test (see below, the yellow points in Figure 4F). We also confirmed the difference between the tVNS and sham conditions using a post-hoc direct comparison on the electrodes marked in blue in Figure 4A, using Wilcoxon signed rank tests (Results).

### Statistical Analyses

Unless stated otherwise, all statistical tests were carried out using a Wilcoxon signed rank test (Wilcoxon, 1945). This included the significance of the pupil time-course, which was corrected for multiple comparisons using FDR correction (Benjamini and Yekutieli, 2011). The significance for alpha attenuation in the EEG spectrogram was assessed using a cluster permutation test with the Monte Carlo method and a dependent samples T-statistic with 10,000 permutations, as implemented in the fieldtrip toolbox (Oostenveld et al., 2011). An alpha of 0.05 was considered significant after FDR correction for clusters (Benjamini and Yekutieli, 2011). In Figure 4, we plotted alpha attenuation at the individual participant level, to facilitate, for the reader, the assessment of effect size across participants; these per-subject values were also tested using the Wilcoxon signed rank test (Figure 4C). Data are expressed as mean ± standard error of the mean throughout.

## Results

To investigate the short-term effects of tVNS in naïve humans, we compared pupil dynamics and EEG alpha oscillations in healthy young male volunteers (n=24) induced by multiple trials of short (3.4s) electrical stimulation at the cymba conchae (tVNS) and at the earlobe (sham) (Figure 1). Stimulation was applied at a frequency of 25Hz, with the intensity ramping up during the trial, to a maximal value selected per participant and location (Methods).

First, we verified that the sham and tVNS conditions did not differ in any of the parameters of subjective averseness examined. Indeed, we did not find any significant differences between the tVNS and sham conditions in subjective reports such as pain and irritation (Table 1, P>0.05 for all comparisons after FDR correction). Regarding objective current intensity, the mean values of the currents applied were 2.20±0.24mA in the tVNS condition and 2.79±0.27mA in the sham condition. The higher current intensity in the sham condition was statistically significant (p=0.0125 Wilcoxon signed rank test). This is likely due to lower sensitivity at the earlobe, and “works against” our a-priori hypothesis (larger effects in tVNS *despite* higher current intensity in sham, see Discussion). This finding shows that earlobe stimulation provides good somatosensory control, which distills the changes related specifically to tVNS.

### Transcutaneous vagus nerve stimulation induces pupil dilation

tVNS led to robust pupil dilation that increased gradually (consistent with the ramping up of the stimulation intensity), reaching half maximum at 2.53s after stimulation onset, peaking at 4.25s after stimulation onset, decreasing back to half maximum at 8.17s, and returning to baseline levels 10s after stimulation. During peak pupil dilation, mean pupil size (in pixels) was 4.05±0.92% above baseline (Figure 2A).

In contrast, sham stimulation led to only modest pupil dilation, mean 1.67+0.63%, and peaking around the same time. This dilation level was significantly weaker than following tVNS (p<0.05 between 2.88-5.96s, repeated Wilcoxon signed rank test across all time points, and FDR correction for multiple comparisons, red bar, Figure 2A). These results were largely consistent across individual participants (Figure 2B) and evident in most (21 of 24) participants (for examples see Figure 2C).

Baseline pupil values did not differ significantly between the tVNS and sham conditions (p = 0.5). In both conditions, the smaller the pupil before a specific trial, the higher the stimulation-evoked response (Pearson’s correlation R=−0.27, p<10^−20^). We found no significant differences between the conditions, in blink rate or gaze position (p>0.6 for all comparisons, using the same statistical procedure). In addition, the higher pupil dilation upon tVNS remained significant and robust across individuals, when the blink data were discarded (rather than interpolated). To verify that the effect of pupil dilation was not mediated by the difference in objective currents, we calculated the correlation between differences in pupil dilation (across the tVNS and sham conditions), and the difference in current (across the tVNS and sham conditions). This did not reveal a significant correlation (Spearman correlation R=−0.12, p=0.56).

In a pilot experiment that employed 30s ON / 30s OFF ‘clinical-like’ stimulation, we observed only a modest trend for greater pupil dilation for tVNS than sham stimulation (p = 0.053, n=23), and pupil size did not differ significantly between ON and OFF periods. Thus, short tVNS pulses lead to significantly greater pupil dilation than following sham stimulation. This indicates that tVNS promotes arousal above and beyond somatosensory stimulation at the ear.

### Transcutaneous vagus nerve stimulation attenuates alpha oscillations

Alpha oscillations exhibit considerable inter-individual variability in frequency and scalp topography (Haegens et al., 2014). To discern the effects of tVNS and sham stimulation on alpha activity, we first identified the frequency and topography of alpha oscillations in each participant separately, using PARAFAC analysis (Harshman, 1970). PARAFAC provides a unique solution for decomposing the EEG signal to three factors (time, frequency, channel; Figure 3A) and may enhance sensitivity. This analytic technique was previously applied to electrophysiological recordings (Miwakeichi et al., 2004; Yanagawa et al., 2013; Meij et al., 2016), and specifically to assessing individual alpha oscillations (Barzegaran et al., 2017; Knyazeva et al., 2018); for a detailed review of its EEG applications, see Cong et al (2015). We identified the regions and frequencies of interest for alpha oscillations in each participant separately, using unbiased “break” data between stimulation blocks (Figure 3B). Figure 3C,D presents the result of this process in representative participants, and Figure 4A shows the median region and frequency profile of alpha oscillations across all participants. PARAFAC successfully identified alpha activity (see the examples in Figure 3C,D), capturing each individual’s specific alpha frequency around 7-13Hz, with the typical occipital topography.

After identifying alpha activity for each individual, we quantified the extent that this activity may be decreased by tVNS or sham stimulation in each participant separately. We found that tVNS attenuated alpha activity (mean: 94.35±2.2% of baseline) to a greater extent (p=0.0027, Wilcoxon signed rank test) than did sham stimulation, which was not associated with significant alpha attenuation (mean: 103.55 ±2.4% of baseline). Baseline alpha was not significantly different between the conditions (p = 0.3 via the Wilcoxon signed rank test). Greater alpha attenuation following tVNS was evident in most (19/21) participants (Figure 4B).

We found a significant negative correlation between the differences in alpha attenuation (tVNS vs. sham conditions) in each individual and the differences in applied current (tVNS vs. sham conditions) (Spearman correlation R=−0.49. p=0.02). Accordingly, participants with stronger sham stimulation current showed less difference in alpha attenuation. Along this line, we repeated the analysis for alpha attenuation while removing one third of the participants with the highest difference in current between the conditions. This analysis revealed a difference in alpha attenuation that was even more significant for the remaining 14 participants, despite the fewer number of participants (p=0.0001, Wilcoxon signed rank test). The implication is that the alpha attenuation we observed upon tVNS constitutes a lower bound (an equivalent objective current intensity in the two conditions leads to a stronger difference in alpha attenuation between the tVNS and sham conditions, see also Discussion).”

tVNS-induced alpha attenuation was not observed in our pilot experiment, which employed 30s ON / 30s OFF ‘clinical-like’ stimulation (p > 0.05, n=15). Neither experiment revealed a significant correlation between alpha attenuation and individual subjective (or objective) scores of stimulation intensities, nor a significant correlation between alpha attenuation and pupil dilation at the individual level (all p≥0.1).

To complement the PARAFAC-based analysis and to better understand the precise time-frequency dynamics and topographical changes of alpha attenuation, we used the weighted alpha topography from the break data as a ‘weighted region of interest’. This reduced the data to two dimensions (time and frequency). Such approach ignored the frequency-of-interest and inspected the induced power changes in the 5-15Hz frequency range for the (weighted) occipital region derived from the PARAFAC decomposition (Figure 4A). In line with the previous results, we found that tVNS significantly attenuated activity in the alpha band (8-12Hz, Figure 4G). Similarly, examining the effects of stimulation on EEG dynamics using cluster based permutation (Maris and Oostenveld, 2007) revealed a significant (p=0.0063, white contour in Figure 4G-H) cluster around 8-12Hz in the seconds following stimulation onset. During this time interval, the mean alpha power was 90.84±2.77% in the tVNS condition, significantly lower than the mean 106.66%±2.70% observed in the sham condition (p<0.0001 in a direct comparison). We also compared the two conditions using the classical alpha frequency range (8-12Hz), during stimulation (0-4s) (means: 94.41±2.15% in the tVNS condition and 105.25±2.41% in the sham condition, p=0.0012 (the Wilcoxon signed rank test for both)).

Finally, we examined the extent that the observed alpha attenuation was specific to occipital electrodes. We inspected the topographical changes in voltage around stimulation relative to baseline (Figure 4F). This analysis was carried out by focusing on the a-priori frequency-band of interest derived from the PARAFAC decomposition (Figure 4C), while ignoring the topography-of-interest derived from the break. We examined topographical effects of stimulation on EEG dynamics using topographical cluster-based permutation (Maris and Oostenveld, 2007). This revealed a significant (p<0.05) cluster over occipital electrodes, which exhibited tVNS attenuation > sham attenuation (yellow points in Figure 4F). The implication is that alpha attenuation was specific to occipital areas (the mean attenuation in yellow electrodes in Figure 4F, during 0:4sec: −3.75μV and −1.78μV in the tVNS and sham conditions, respectively, p=0.007 via the Wilcoxon signed rank test). We also compared the two conditions directly using the occipital electrodes marked in Figure 4A, during 0:4sec (mean: 3.75μV and −1.78μV in the respective conditions, p=0.007 via the Wilcoxon signed rank test). Importantly, the regions showing tVNS-induced alpha attenuation overlapped electrodes showing alpha activity in the independent “break” intervals between stimulation blocks (compare the blue dots in Figure 4A with the yellow dots in Figure 4F).

Altogether, the EEG data establish that short tVNS pulses, but not sham stimulation, attenuate occipital alpha activity.

## Discussion

We examined the effects of short tVNS pulses (and sham stimulation at the ear lobe) on pupil dynamics and EEG alpha activity in naïve healthy men. While subjective stimulation intensities were not significantly different in the two conditions (Table 1), we found that short tVNS pulses induce pupil dilation (Figure 2) and EEG alpha attenuation (Figure 4) to a greater extent than does sham stimulation. These effects support the hypothesis that tVNS activates endogenous arousal-promoting neuromodulatory signaling such as LC-NE activity, as is known to occur in invasive VNS (Hulsey et al., 2017; Mridha et al., 2019). This suggests that tVNS mimics VNS.

### Validity and limitations

Our results were obtained during fixation at rest. Although they may be applicable to other conditions, future studies are needed to determine the effects of short-pulse tVNS during other states, such as drowsiness and sleep, and during the performance of specific cognitive tasks. For example, high arousal at baseline could create a ceiling effect for pupil dilation and alpha attenuation; conversely, during decreased vigilance, EEG effects may attenuate idle activity at different frequency bands (e.g. changing the theta/alpha ratio during drowsiness, or suppressing slow wave activity in sleep). Another limitation is that we could only study tVNS in male volunteers. Since there may be sex-specific differences in LC-NE and neuromodulatory activity (Bangasser et al., 2016), future studies with females are warranted. Lastly, our experimental design equated the subjective intensity of tVNS and sham stimulation. Our data revealed that a significantly higher current at the earlobe (sham condition) was necessary to achieve equal subjective intensity. We did not find a significant correlation between differences in current (tVNS vs. sham) and in pupil dilation (tVNS vs. sham), but current differences were negatively correlated with differences in alpha attenuation (tVNS vs. sham). Accordingly, among participants with a higher current intensity in the sham than the tVNS condition, the difference in alpha attenuation was smaller between the conditions. Indeed, restricting the analysis to a subset of 14 individuals, such that the significant difference in current intensity was eliminated, revealed a stronger effect of alpha attenuation in the tVNS vs. sham condition. Thus, our results represent a conservative lower-bound of the actual difference between alpha attenuation in tVNS and sham, which would be even greater in the context of comparable currents in tVNS and sham. More generally, this issue is relevant to an inherent limitation of using earlobe sham stimulation as a control condition. Despite its extensive use (Yap et al., 2020) and advantages, the earlobe, with its lower sensitivity, requires higher currents to produce a comparable subjective intensity. Future studies should apply additional control conditions (e.g. stimulation at other frequencies) to mitigate this limitation.

### Previous tVNS studies

Our finding that tVNS attenuates alpha oscillations is compatible with the findings of a number of studies (Bodin et al., 2015; Lewine et al., 2019); while earlier studies reported mixed results or did not detect EEG effects (Hammond et al., 1992; Salinsky and Burchiel, 1993). Our use of short tVNS pulses likely contributed to our ability to reveal alpha attenuation. In addition, the sensitive analysis that was afforded by the use of PARAFAC enabled identifying alpha effects in many, but not all the participants.

In contrast to our focus on ongoing EEG and pupillometry, most previous studies attempted to demonstrate the effectiveness of tVNS by focusing on the EEG P300 or on salivary alpha amylase as readouts. The P300 is a positive deflection with maximal amplitude in electrodes placed over the centro-parietal midline, 300-500ms after stimulus onset. The amplitude of this deflection is modulated by the probability of stimulus appearance regardless of sensory modality (Desmedt et al., 1965; Sutton et al., 1965). The P300 has been hypothesized to be a marker of LC-NE activity (Nieuwenhuis et al., 2005). This is because LC neurons are likewise activated by infrequent stimuli, independent of sensory modality (Aston-Jones et al., 1991); and deviant stimuli elicit greater pupil dilation than standard stimuli (Murphy et al., 2011). However, the P300 may not constitute a straightforward test of tVNS efficacy since the link between P300 and LC-NE activity is still debated (Nieuwenhuis et al., 2011), and the dopaminergic (Glover et al., 1988) and glutamatergic (Hall et al., 2015) systems could also substantially affect the P300. Ventura-Bort et al (2018) demonstrated that tVNS amplifies the parietal component of the P300 effect (P3b), selectively, for easy targets in their task. However, this effect was modest and could not be replicated using weaker fixed currents (0.5mA) and a simpler classical P300 task (Warren et al., 2018). Another study by Keute et al (2019) focused on the difference in pupil dilation between deviant and standard stimuli, using a classical auditory oddball task. The use of a constant 3mA tVNS in all participants did not reveal any effect of the stimulation on event-related or baseline pupil size. A possible explanation is that 30s tVNS modulates tonic NE levels but does not affect phasic stimulus-evoked changes in NE that are associated with the P300. In agreement with this possibility, the use of clonidine (an α2 adrenergic receptor agonist that reduces NE signaling) provided similar mixed results (Pineda and Swick, 1992; Halliday et al., 1994; Pineda et al., 1997; Brown et al., 2015). Future studies that will use short tVNS pulses, as used here, may help elucidate the effects on the P300.

Both Warren et al (Warren et al., 2018) and Ventura-Bort (2018) showed that tVNS increases levels of salivary alpha amylase, which has served as a peripheral measure of sympathetic activity associated with LC-NE signaling (Rohleder and Nater, 2009). However, this measure has poor temporal resolution and can only reveal differences between time intervals before vs. after stimulation blocks that last several minutes. This approach does not leverage the superior temporal resolution of tVNS compared to pharmacological manipulations of NE in humans; such manipulations are highly effective in studying the effects of slower NE dynamics (Gelbard-Sagiv et al., 2018). By contrast, the transient (within seconds) tVNS-mediated effects revealed here offer considerable advantages over the slow modulations elicited by NE drugs.

### Pupil dilation and alpha attenuation as indices of arousal and LC-NE activity

Pupil diameter was suggested as a proxy for noradrenergic signaling since Aston Jones and Cohen first provided an example of correlated dynamics in simultaneous pupil and LC single-unit activities in a monkey (Aston-Jones and Cohen, 2005; and see a recent review by Joshi and Gold, 2020). Since this initial report, the relation between pupil diameter and noradrenergic signaling has been established in monkeys (Varazzani et al., 2015; Joshi et al., 2016), rats (Liu et al., 2017; Hayat et al., 2020), and mice (Reimer et al., 2016; Breton-Provencher and Sur, 2019), as well as in human BOLD fMRI (Murphy et al., 2014). The tVNS-induced pupil dilation time-course that we observed (Figure 3) resembles pupil dynamics in response to LC electrical stimulation in monkeys (Joshi et al., 2016) and optogenetic stimulation in rats (Hayat et al., 2020). This supports the hypothesis that tVNS activates the LC, as has been established for invasive VNS.

Alpha oscillations are abundant during detachment from the sensory environment in wakefulness. These are considered an index of low arousal (Torsvall and Akerstedt, 1987b; Drapeau and Carrier, 2004b). Alpha oscillations are believed to represent an “idling” state of cortical activity (Steriade, 2001; Palva and Palva, 2007) that is expected to be anti-correlated with arousal-promoting activity, such as that of the LC-NE system. These oscillations bias sensory perception (Waschke et al., 2019). A recent study that used long 2min tVNS on the neck also found that tVNS attenuates alpha and theta oscillation (Lewine et al., 2019).

### LC-NE vs. other neuromodulatory systems

While pupil dilation and EEG alpha attenuation are both *compatible* with noradrenergic signaling, LC-NE involvement is unlikely to be the only modulatory system involved, given the overlap and redundancy among neuromodulatory systems. Other elements such as the cholinergic system also contribute to brain arousal and are associated with both pupil dilation (Reimer et al., 2016) and EEG activation (Szerb, 1967). However, cholinergic activation alone is unlikely to drive the effects observed. This is because during rapid eye movement (REM) sleep, cholinergic activation occurs without LC-NE activity (Nir and Tononi, 2010); and the EEG is activated but pupils remain constricted (Siegel, 2005). Moreover, given that VNS robustly activates the LC, and no such relation has been reported for cholinergic nuclei, the most parsimonious interpretation is that the primary neuromodulatory effects of tVNS are noradrenergic, while cholinergic modulation (Mridha et al., 2019) is likely secondary. tVNS may engage additional subcortical neuromodulatory systems such as the dorsal raphe and the ventral tegmental area, as observed with tVNS-induced BOLD fMRI (Frangos et al., 2015). Thus, the possible relation of tVNS to other neuromodulatory systems beyond LC-NE is an important topic for further investigation.

### tVNS as a novel tool for transient neuromodulation

Great interest has arisen in investigating the contribution of the LC-NE system to human cognition, including perception, learning and memory, decision-making, and aging and neurodegeneration. In this context, tVNS entails important advantages over existing tools. While important advances have been made by relying on the correlation of LC-NE activity with pupil dynamics (e.g de Gee et al., 2017), hidden factors (such as fluctuations in arousal and attention) could be at the basis of the observed correlations (Clewett et al., 2018; Dragone et al., 2018). Previous human studies also employed *causal* perturbations, using NE drugs to study effects on perception (Gelbard-Sagiv et al., 2018), memory (see van Stegeren, 2008 for review), and decision making (Warren et al., 2017). However, the systemic delivery of NE drugs is inherently limited to affecting tonic LC-NE activity and has poor temporal resolution, whereas tVNS has clear added value.

A number of studies reported a benefit of invasive VNS on memory (Clark et al., 1999; Jacobs et al., 2015; for review see Hansen, 2017; Sun et al., 2017). However, the participants of those studies had severe epilepsy or depression. In addition, ongoing daily VNS induces complex long-term plastic changes that make interpretation difficult.

Due to limitations of the available techniques for studying cognition, the potential of tVNS has been recognized and is increasingly being realized (Van Leusden et al., 2015). However, to date, the evidence supporting the effectiveness of tVNS in mimicking invasive VNS is mixed. We applied short stimulation pulses and limited currents to a maximal value per participant. Accordingly, and by focusing on simple pupillary and ongoing EEG readouts, we showed that tVNS transiently elicits markers of brain arousal that are compatible with arousal-promoting neuromodulatory signaling such as NE/Ach. This supports the hypothesis that tVNS mimics invasive VNS, thereby extending the experimental toolkit for non-pharmacological neuromodulation in humans with high temporal resolution. Therefore, tVNS can be used to further investigate the means by which transient neuromodulation contributes to human cognition. Future studies should compare several levels of both sham stimulation and tVNS, to conduct a parametric investigation. Of note, stronger stimulation may not necessarily produce stronger effects on neuronal activity and behavior, since some effects may actually elicit “U-shape” profiles (for example see Clark et al., 1999).

Finally, tVNS should be conducted to further elucidate the processes mediating the clinical improvements elicited by VNS in epilepsy and depression. These include the role of arousal-promoting neuromodulatory signaling in improving mood in depressed patients (Grimonprez et al., 2015; Liu et al., 2016; Fang et al., 2017; Tu et al., 2018). In particular, tVNS-induced pupillary and EEG effects may help predict clinical efficacy of invasive VNS and thus facilitate triaging patients to receive either conservative therapy or surgical implantation of VNS stimulation devices.

## Software Accessibility

code is available upon request.

## Acknowledgements

Supported by ISF 51/11 (I-CORE cognitive sciences) and the Adelis Foundation (YN), The Herczeg Institute on Aging, the TAU global research fund, and the Naomi Foundation (OS). The authors thank members of the Nir lab for discussions, and Leon Deouell and Amit Marmelshtein for their comments on earlier drafts of the manuscript.

## References

Amzica F, Lopes da Silva FH (2017) Niedermeyer’s Electroencephalography (Schomer DL, Lopes da Silva FH, eds). Oxford University Press. Available at: http://www.oxfordmedicine.com/view/10.1093/med/9780190228484.001.0001/med-9780190228484.

Andersson CA, Bro R (2000) The N-way Toolbox for MATLAB. 3–6.

Aston-Jones G, Chiang C, Alexinsky T (1991) Discharge of noradrenergic locus coeruleus neurons in behaving rats and monkeys suggests a role in vigilance. Prog Brain Res 88:501–520 Available at: http://www.ncbi.nlm.nih.gov/pubmed/1813931.

Aston-Jones G, Cohen JD (2005) An integrative theory of locus coeruleus-norepinephrine function: adaptive gain and optimal performance. Annu Rev Neurosci 28:403–450 Available at: http://www.ncbi.nlm.nih.gov/pubmed/16022602.

Badran BW, Brown JC, Dowdle LT, Mithoefer OJ, LaBate NT, Coatsworth J, DeVries WH, Austelle CW, McTeague LM, Yu A, Bikson M, Jenkins DD, George MS (2018) Tragus or cymba conchae? Investigating the anatomical foundation of transcutaneous auricular vagus nerve stimulation (taVNS). Brain Stimul 11:947–948 Available at: https://doi.org/10.1016/j.brs.2018.06.003.

Bangasser DA, Wiersielis KR, Khantsis S (2016) Sex differences in the locus coeruleus-norepinephrine system and its regulation by stress. Brain Res 1641:177–188 Available at: http://www.ncbi.nlm.nih.gov/pubmed/26607253.

Barzegaran E, Vildavski VY, Knyazeva MG (2017) Fine Structure of Posterior Alpha Rhythm in Human EEG : Frequency Components, Their Cortical Sources, and Temporal Behavior. 1–12.

Bauer S, Baier H, Baumgartner C, Bohlmann K, Fauser S, Graf W, Hillenbrand B, Hirsch M, Last C, Lerche H, Mayer T, Schulze-Bonhage A, Steinhoff BJ, Weber Y, Hartlep A, Rosenow F, Hamer HM (2016) Transcutaneous Vagus Nerve Stimulation (tVNS) for Treatment of Drug-Resistant Epilepsy: A Randomized, Double-Blind Clinical Trial (cMPsE02). Brain Stimul 9:356–363 Available at: http://dx.doi.org/10.1016/j.brs.2015.11.003.

Benjamini Y, Yekutieli D (2011) The Control of the False Discovery Rate in Multiple Testing under Dependency Source. Statistics (Ber) 29:1165–1188.

Bermejo P, López M, Larraya I, Chamorro J, Cobo JL, Ordóñez S, Vega JA (2017) Innervation of the Human Cavum Conchae and Auditory Canal : Anatomical Basis for Transcutaneous Auricular Nerve Stimulation. 2017.

Bodin C, Aubert S, Daquin G, Carron R, Scavarda D, McGonigal A, Bartolomei F (2015) Responders to vagus nerve stimulation (VNS) in refractory epilepsy have reduced interictal cortical synchronicity on scalp EEG. Epilepsy Res 113:98–103 Available at: http://dx.doi.org/10.1016/j.eplepsyres.2015.03.018.

Boon P, De Cock E, Mertens A, Trinka E (2018) Neurostimulation for drug-resistant epilepsy. Curr Opin Neurol 31:198–210 Available at: http://insights.ovid.com/crossref?an=00019052-201804000-00015.

Breton-Provencher V, Sur M (2019) Active control of arousal by a locus coeruleus GABAergic circuit. Nat Neurosci 22:218–228 Available at: http://www.ncbi.nlm.nih.gov/pubmed/30643295.

Bro R, Kiers HAL (2003) A new efficient method for determining the number of components in PARAFAC models. J Chemom 17:274–286 Available at: http://doi.wiley.com/10.1002/cem.801.

Brown SBRE, van der Wee NJA, van Noorden MS, Giltay EJ, Nieuwenhuis S (2015) Noradrenergic and cholinergic modulation of late ERP responses to deviant stimuli. Psychophysiology 52:1620–1631 Available at: http://doi.wiley.com/10.1111/psyp.12544.

Clark KB, Naritoku DK, Smith DC, Browning RA, Jensen RA (1999) Enhanced recognition memory following vagus nerve stimulation in human subjects. Nat Neurosci 2:94–98 Available at: http://www.nature.com/doifinder/10.1038/4600.

Clewett D V., Huang R, Velasco R, Lee T-H, Mather M (2018) Locus Coeruleus Activity Strengthens Prioritized Memories Under Arousal. J Neurosci 38:1558–1574 Available at: http://www.jneurosci.org/lookup/doi/10.1523/JNEUROSCI.2097-17.2017.

Cong F, Lin Q, Kuang L, Gong X, Astikainen P (2015) Tensor decomposition of EEG signals : A brief review. J Neurosci Methods 248:59–69 Available at: http://dx.doi.org/10.1016/j.jneumeth.2015.03.018.

de Gee JW, Colizoli O, Kloosterman NA, Knapen T, Nieuwenhuis S, Donner TH (2017) Dynamic modulation of decision biases by brainstem arousal systems. Elife 6 Available at: http://www.ncbi.nlm.nih.gov/pubmed/28383284.

Desbeaumes Jodoin V, Lespérance P, Nguyen DK, Fournier-Gosselin M-PP, Richer F (2015) Effects of vagus nerve stimulation on pupillary function. Int J Psychophysiol 98:455–459 Available at: http://dx.doi.org/10.1016/j.ijpsycho.2015.10.001.

Desmedt JE, Debecker J, Manil J (1965) [Demonstration of a cerebral electric sign associated with the detection by the subject of a tactile sensorial stimulus. The analysis of cerebral evoked potentials derived from the scalp with the aid of numerical ordinates]. Bull Acad R Med Belg 5:887–936 Available at: http://www.ncbi.nlm.nih.gov/pubmed/5864251.

Dorr AE, Debonnel G (2006) Effect of Vagus Nerve Stimulation on Serotonergic and Noradrenergic Transmission. 318:890–898.

Dragone A, Lasaponara S, Pinto M, Rotondaro F, De Luca M, Doricchi F (2018) Expectancy modulates pupil size during endogenous orienting of spatial attention. Cortex 102:57–66 Available at: https://linkinghub.elsevier.com/retrieve/pii/S0010945217303179.

Drapeau C, Carrier J (2004a) Fluctuation of waking electroencephalogram and subjective alertness during a 25-hour sleep-deprivation episode in young and middle-aged subjects. Sleep 27:55–60 Available at: http://www.ncbi.nlm.nih.gov/pubmed/14998238.

Drapeau C, Carrier J (2004b) Fluctuation of waking electroencephalogram and subjective alertness during a 25-hour sleep-deprivation episode in young and middle-aged subjects. Sleep 27:55–60.

Ellrich J (2011) Transcutaneous Vagus Nerve Stimulation. Eur Neurol Rev 6:254 Available at: http://www.touchneurology.com/articles/transcutaneous-vagus-nerve-stimulation.

Fang J, Egorova N, Rong P, Liu J, Hong Y, Fan Y, Wang X, Wang H, Yu Y, Ma Y, Xu C, Li S, Zhao J, Luo M, Zhu B, Kong J (2017) Early cortical biomarkers of longitudinal transcutaneous vagus nerve stimulation treatment success in depression. NeuroImage Clin 14:105–111 Available at: http://dx.doi.org/10.1016/j.nicl.2016.12.016.

Follesa P, Biggio F, Gorini G, Caria S, Talani G, Dazzi L, Puligheddu M, Marrosu F, Biggio G (2007) Vagus nerve stimulation increases norepinephrine concentration and the gene expression of BDNF and bFGF in the rat brain. 79.

Frangos E, Ellrich J, Komisaruk BR (2015) Non-invasive access to the vagus nerve central projections via electrical stimulation of the external ear: FMRI evidence in humans. Brain Stimul 8:624–636 Available at: http://dx.doi.org/10.1016/j.brs.2014.11.018.

Gelbard-Sagiv H, Magidov E, Sharon H, Hendler T, Nir Y (2018) Noradrenaline Modulates Visual Perception and Late Visually Evoked Activity. Curr Biol 28:2239–2249.e6 Available at: http://www.ncbi.nlm.nih.gov/pubmed/29983318.

Glover A, Ghilardi MF, Bodis-Wollner I, Onofrj M (1988) Alterations in event-related potentials (ERPs) of MPTP-treated monkeys. Electroencephalogr Clin Neurophysiol Potentials Sect 71:461–468 Available at: https://linkinghub.elsevier.com/retrieve/pii/0168559788900500.

Grimonprez A et al. (2015) The antidepressant-like effect of vagus nerve stimulation is mediated through the locus coeruleus. J Psychiatr Res 68:1–7 Available at: http://dx.doi.org/10.1016/j.jpsychires.2015.05.002.

Groves DA, Bowman EM, Brown VJ (2005) Recordings from the rat locus coeruleus during acute vagal nerve stimulation in the anaesthetised rat. Neurosci Lett 379:174–179.

Haegens S, Cousijn H, Wallis G, Harrison PJ, Nobre AC (2014) Inter- and intra-individual variability in alpha peak frequency. Neuroimage 92:46–55 Available at: http://dx.doi.org/10.1016/j.neuroimage.2014.01.049.

Hall M-H, Jensen JE, Du F, Smoller JW, O’Connor L, Spencer KM, Öngür D (2015) Frontal P3 event-related potential is related to brain glutamine/glutamate ratio measured in vivo. Neuroimage 111:186–191 Available at: https://linkinghub.elsevier.com/retrieve/pii/S1053811915001159.

Halliday R, Naylor H, Brandeis D, Callaway E, Yano L, Herzig K (1994) The effect of D-amphetamine, clonidine, and yohimbine on human information processing. Psychophysiology 31:331–337.

Hammond EJ, Uthman BM, Reid SA, Wilder BJ (1992) Electrophysiological Studies of Cervical Vagus Nerve Stimulation in Humans: I. EEG Effects. Epilepsia 33:1013–1020.

Hansen N (2017) The Longevity of Hippocampus-Dependent Memory Is Orchestrated by the Locus Coeruleus-Noradrenergic System. Neural Plast 2017.

Harshman RA (1970) Foundations of the PARAFAC procedure: Models and conditions for an “explanatory” multimodal factor analysis.

Hayat H, Regev N, Matosevich N, Sales A, Paredes-Rodriguez E, Krom AJ, Bergman L, Li Y, Lavigne M, Kremer EJ, Yizhar O, Pickering AE, Nir Y (2020) Locus coeruleus norepinephrine activity mediates sensory-evoked awakenings from sleep. Sci Adv 6:eaaz4232 Available at: http://www.ncbi.nlm.nih.gov/pubmed/32285002.

He W, Jing X-H, Zhu B, Zhu X-L, Li L, Bai W-Z, Ben H (2013) The auriculo-vagal afferent pathway and its role in seizure suppression in rats. BMC Neurosci 14:85 Available at: http://bmcneurosci.biomedcentral.com/articles/10.1186/1471-2202-14-85.

Hein E, Nowak M, Kiess O, Biermann T, Bayerlein K, Kornhuber J, Kraus T (2013) Auricular transcutaneous electrical nerve stimulation in depressed patients: A randomized controlled pilot study. J Neural Transm 120:821–827.

Hulsey DR, Riley JR, Loerwald KW, Ii RLR, Kilgard MP, Hays SA (2017) Parametric characterization of neural activity in the locus coeruleus in response to vagus nerve stimulation. Exp Neurol 289:21–30 Available at: http://dx.doi.org/10.1016/j.expneurol.2016.12.005.

Jacobs HIL, Riphagen JM, Razat CM, Wiese S, Sack AT (2015) Transcutaneous vagus nerve stimulation boosts associative memory in older individuals. Neurobiol Aging 36:1860–1867.

Joshi S, Gold JI (2020) Pupil Size as a Window on Neural Substrates of Cognition. Trends Cogn Sci 24:466–480 Available at: https://linkinghub.elsevier.com/retrieve/pii/S1364661320300802.

Joshi S, Li Y, Kalwani RM, Gold JI (2016) Relationships between Pupil Diameter and Neuronal Activity in the Locus Coeruleus, Colliculi, and Cingulate Cortex. Neuron 89:221–234 Available at: https://linkinghub.elsevier.com/retrieve/pii/S089662731501034X.

Keute M, Demirezen M, Graf A, Mueller NG, Zaehle T (2019) No modulation of pupil size and event-related pupil response by transcutaneous auricular vagus nerve stimulation (taVNS). Sci Rep 9:11452 Available at: http://www.nature.com/articles/s41598-019-47961-4.

Knyazeva MG, Barzegaran E, Vildavski VY, Demonet J (2018) Neurobiology of Aging of human alpha rhythm. Neurobiol Aging 69:261–273 Available at: https://doi.org/10.1016/j.neurobiolaging.2018.05.018.

Krahl SE, Clark KB (2012) Vagus nerve stimulation for epilepsy: A review of central mechanisms. Surg Neurol Int 3:S255–9 Available at: http://www.ncbi.nlm.nih.gov/pubmed/23230530.

Krahl SE, Clark KB, Smith DC, Browning RA (1998) Locus coeruleus lesions suppress the seizure-attenuating effects of vagus nerve stimulation. Epilepsia 39:709–714.

Kraus T, Hösl K, Kiess O, Schanze A, Kornhuber J, Forster C (2007) BOLD fMRI deactivation of limbic and temporal brain structures and mood enhancing effect by transcutaneous vagus nerve stimulation. J Neural Transm 114:1485–1493.

Kraus T, Kiess O, Hösl K, Terekhin P, Kornhuber J, Forster C (2013) CNS BOLD fMRI effects of sham-controlled transcutaneous electrical nerve stimulation in the left outer auditory canal - A pilot study. Brain Stimul 6:798–804 Available at: http://dx.doi.org/10.1016/j.brs.2013.01.011.

Kwon C-S, Ripa V, Al-Awar O, Panov F, Ghatan S, Jetté N (2018) Epilepsy and Neuromodulation—Randomized Controlled Trials. Brain Sci 8:69 Available at: http://www.mdpi.com/2076-3425/8/4/69.

Lewine JD, Paulson K, Bangera N, Simon BJ (2019) Exploration of the Impact of Brief Noninvasive Vagal Nerve Stimulation on EEG and Event-Related Potentials. Neuromodulation Technol Neural Interface 22:564–572 Available at: https://onlinelibrary.wiley.com/doi/abs/10.1111/ner.12864.

Liu J, Fang J, Wang Z, Rong P, Hong Y, Fan Y, Wang X, Park J, Jin Y, Liu C, Zhu B, Kong J (2016) Transcutaneous vagus nerve stimulation modulates amygdala functional connectivity in patients with depression. J Affect Disord 205:319–326 Available at: http://dx.doi.org/10.1016/j.jad.2016.08.003.

Liu Y, Rodenkirch C, Moskowitz N, Schriver B, Wang Q (2017) Dynamic Lateralization of Pupil Dilation Evoked by Locus Coeruleus Activation Results from Sympathetic, Not Parasympathetic, Contributions. Cell Rep 20:3099–3112 Available at: http://www.ncbi.nlm.nih.gov/pubmed/28954227.

Manta S, Dong J, Debonnel G, Blier P (2009) Enhancement of the function of rat serotonin and norepinephrine neurons by sustained vagus nerve stimulation. J Psychiatry Neurosci 34:272–280.

Manta S, Mansari M El, Debonnel G, Blier P (2013) Electrophysiological and neurochemical effects of long-term vagus nerve stimulation on the rat monoaminergic systems. 459–470.

Maris E, Oostenveld R (2007) Nonparametric statistical testing of EEG− and MEG-data. J Neurosci Methods 164:177–190.

Meij R Van Der, Ede F Van, Maris E (2016) Rhythmic Components in Extracranial Brain Signals Reveal Multifaceted Task Modulation of Overlapping Neuronal Activity. 1–28.

Miwakeichi F, Martínez-Montes E, Valdés-Sosa PA, Nishiyama N, Mizuhara H, Yamaguchi Y (2004) Decomposing EEG data into space-time-frequency components using Parallel Factor Analysis. Neuroimage 22:1035–1045 Available at: http://www.ncbi.nlm.nih.gov/pubmed/15219576.

Mridha Z, Gee JW de, Shi Y, Alkashgari R, Williams J, Suminski A, Ward MP, Zhang W, McGinley MJ (2019) Graded recruitment of pupil-linked neuromodulation by parametric stimulation of the vagus nerve. bioRxiv:2019.12.28.890111 Available at: http://biorxiv.org/content/early/2019/12/30/2019.12.28.890111.abstract.

Murphy PR, O’Connell RG, O’Sullivan M, Robertson IH, Balsters JH (2014) Pupil diameter covaries with BOLD activity in human locus coeruleus. Hum Brain Mapp 35:4140–4154 Available at: http://www.ncbi.nlm.nih.gov/pubmed/24510607.

Murphy PR, Robertson IH, Balsters JH, O’connell RG (2011) Pupillometry and P3 index the locus coeruleus-noradrenergic arousal function in humans. Psychophysiology 48:1532–1543.

Nemeroff CB, Mayberg HS, Krahl SE, McNamara J, Frazer A, Henry TR, George MS, Charney DS, Brannan SK (2006) VNS therapy in treatment-resistant depression: clinical evidence and putative neurobiological mechanisms. Neuropsychopharmacology 31:1345–1355 Available at: http://www.ncbi.nlm.nih.gov/pubmed/16641939.

Nieuwenhuis S, Aston-Jones G, Cohen JD (2005) Decision making, the P3, and the locus coeruleus-norepinephrine system. Psychol Bull 131:510–532 Available at: http://doi.apa.org/getdoi.cfm?doi=10.1037/0033-2909.131.4.510.

Nieuwenhuis S, De Geus EJ, Aston-Jones G (2011) The anatomical and functional relationship between the P3 and autonomic components of the orienting response. Psychophysiology 48:162–175 Available at: http://www.ncbi.nlm.nih.gov/pubmed/20557480.

Nir Y, Tononi G (2010) Dreaming and the brain: from phenomenology to neurophysiology. Trends Cogn Sci 14:88–100 Available at: http://www.pubmedcentral.nih.gov/articlerender.fcgi?artid=2814941&tool=pmcentrez&rendertype=abstract [Accessed May 28, 2014].

Oostenveld R, Fries P, Maris E, Schoffelen J-M (2011) FieldTrip: Open Source Software for Advanced Analysis of MEG, EEG, and Invasive Electrophysiological Data. Comput Intell Neurosci 2011:1–9.

Palva S, Palva JM (2007) New vistas for α-frequency band oscillations. Trends Neurosci 30:150–158.

Peirce JW (2007) PsychoPy--Psychophysics software in Python. J Neurosci Methods 162:8–13 Available at: http://www.ncbi.nlm.nih.gov/pubmed/17254636.

Peuker ET, Filler TJ (2002) The nerve supply of the human auricle. Clin Anat 15:35–37.

Pineda JA, Swick D (1992) Visual P3-like potentials in squirrel monkey: Effects of a noradrenergic agonist. Brain Res Bull 28:485–491 Available at: https://linkinghub.elsevier.com/retrieve/pii/036192309290051X.

Pineda JA, Westerfield M, Kronenberg BM, Kubrin J (1997) Human and monkey P3-like responses in a mixed modality paradigm: effects of context and context-dependent noradrenergic influences. Int J Psychophysiol 27:223–240 Available at: https://linkinghub.elsevier.com/retrieve/pii/S0167876097000615.

Raedt R, Clinckers R, Mollet L, Vonck K, El Tahry R, Wyckhuys T, De Herdt V, Carrette E, Wadman W, Michotte Y, Smolders I, Boon P, Meurs A (2011) Increased hippocampal noradrenaline is a biomarker for efficacy of vagus nerve stimulation in a limbic seizure model. J Neurochem 117:461–469 Available at: http://doi.wiley.com/10.1111/j.1471-4159.2011.07214.x.

Redgrave J, Day D, Leung H, Laud PJ, Ali A, Lindert R, Majid A (2018) Safety and tolerability of Transcutaneous Vagus Nerve stimulation in humans; a systematic review. Brain Stimul 11:1225–1238 Available at: https://doi.org/10.1016/j.brs.2018.08.010.

Reimer J, McGinley MJ, Liu Y, Rodenkirch C, Wang Q, McCormick DA, Tolias AS (2016) Pupil fluctuations track rapid changes in adrenergic and cholinergic activity in cortex. Nat Commun 7:13289 Available at: http://www.nature.com/articles/ncomms13289.

Rohleder N, Nater UM (2009) Determinants of salivary alpha-amylase in humans and methodological considerations. Psychoneuroendocrinology 34:469–485 Available at: http://www.ncbi.nlm.nih.gov/pubmed/19155141.

Rong P, Liu J, Wang L, Liu R, Fang J, Zhao J, Zhao Y, Wang H, Vangel M, Sun S, Ben H, Park J, Li S, Meng H, Zhu B, Kong J (2016) Effect of transcutaneous auricular vagus nerve stimulation on major depressive disorder: A nonrandomized controlled pilot study. J Affect Disord 195:172–179 Available at: http://dx.doi.org/10.1016/j.jad.2016.02.031.

Roosevelt RW, Smith DC, Clough RW, Jensen RA, Browning RA (2006) Increased extracellular concentrations of norepinephrine in cortex and hippocampus following vagus nerve stimulation in the rat. Brain Res 1119:124–132.

Safi S, Ellrich J, Neuhuber W (2016) Myelinated Axons in the Auricular Branch of the Human Vagus Nerve. Anat Rec 299:1184–1191.

Salinsky MC, Burchiel KJ (1993) Vagus Nerve Stimulation Has No Effect on Awake EEG Rhythms in Humans. Epilepsia 34:299–304.

Sellaro R, van Leusden JWR, Tona K-D, Verkuil B, Nieuwenhuis S, Colzato LS (2015) Transcutaneous Vagus Nerve Stimulation Enhances Post-error Slowing. J Cogn Neurosci 27:2126–2132 Available at: http://www.mitpressjournals.org/doi/10.1162/jocn_a_00851.

Siegel JM (2005) Chapter 10 - REM Sleep. In: Principles And Practice Of Sleep Medicine (Fourth Edition), Fourth Edi. (Kryger MH, Roth T, Dement WC, eds), pp 120–135. Philadelphia: W.B. Saunders. Available at: http://www.sciencedirect.com/science/article/pii/B0721607977500173.

Steenbergen L, Sellaro R, Stock AK, Verkuil B, Beste C, Colzato LS (2015) Transcutaneous vagus nerve stimulation (tVNS) enhances response selection during action cascading processes. Eur Neuropsychopharmacol 25:773–778 Available at: http://dx.doi.org/10.1016/j.euroneuro.2015.03.015.

Stefan H, Kreiselmeyer G, Kerling F, Kurzbuch K, Rauch C, Heers M, Kasper BS, Hammen T, Rzonsa M, Pauli E, Ellrich J, Graf W, Hopfengärtner R (2012) Transcutaneous vagus nerve stimulation (t-VNS) in pharmacoresistant epilepsies: a proof of concept trial. Epilepsia 53:e115–8 Available at: http://www.ncbi.nlm.nih.gov/pubmed/22554199.

Steriade M (2001) Impact of network activities on neuronal properties in corticothalamic systems. J Neurophysiol 86:1–39 Available at: http://www.ncbi.nlm.nih.gov/pubmed/11431485.

Sun L, Peräkylä J, Holm K, Haapasalo J, Lehtimäki K, Ogawa KH, Peltola J, Hartikainen KM (2017) Vagus nerve stimulation improves working memory performance. J Clin Exp Neuropsychol 00:1–11 Available at: https://www.tandfonline.com/doi/full/10.1080/13803395.2017.1285869.

Sutton S, Braren M, Zubin J, John ER (1965) Evoked-potential correlates of stimulus uncertainty. Science 150:1187–1188 Available at: http://www.ncbi.nlm.nih.gov/pubmed/5852977.

Szerb JC (1967) Cortical acetylcholine release and electroencephalographic arousal. J Physiol 192:329–343 Available at: http://www.ncbi.nlm.nih.gov/pubmed/6050151.

Takigawa M, Mogenson GJ (1977) A study of inputs to antidromically identified neurons of the locus coeruleus. Brain Res 135:217–230 Available at: http://www.ncbi.nlm.nih.gov/pubmed/922473.

Torsvall L, Akerstedt T (1987a) Sleepiness on the job: continuously measured EEG changes in train drivers. Electroencephalogr Clin Neurophysiol 66:502–511 Available at: http://www.ncbi.nlm.nih.gov/pubmed/2438115.

Torsvall L, Akerstedt T (1987b) Sleepiness on the job: continuously measured EEG changes in train drivers. Electroencephalogr Clin Neurophysiol 66:502–511.

Trevizol AP, Shiozawa P, Taiar I, Soares A, Gomes JS, Barros MD, Liquidato BM, Cordeiro Q (2016) Transcutaneous Vagus Nerve Stimulation (taVNS) for Major Depressive Disorder: An Open Label Proof-of-Concept Trial. Brain Stimul 9:453–454 Available at: http://dx.doi.org/10.1016/j.brs.2016.02.001.

Tu Y, Fang J, Cao J, Wang Z, Park J, Jorgenson K, Lang C, Liu J, Zhang G, Zhao Y, Zhu B, Rong P, Kong J (2018) A distinct biomarker of continuous transcutaneous vagus nerve stimulation treatment in major depressive disorder. Brain Stimul 11:501–508 Available at: https://doi.org/10.1016/j.brs.2018.01.006.

Van Bockstaele EJ, Peoples J, Telegan P (1999) Efferent projections of the nucleus of the solitary tract to peri-Locus coeruleus dendrites in rat brain: Evidence for a monosynaptic pathway. J Comp Neurol 412:410–428.

Van Leusden JWR, Sellaro R, Colzato LS (2015) Transcutaneous Vagal Nerve Stimulation (tVNS): A new neuromodulation tool in healthy humans? Front Psychol 6:2013–2016.

van Stegeren AH (2008) The role of the noradrenergic system in emotional memory. Acta Psychol (Amst) 127:532–541 Available at: https://linkinghub.elsevier.com/retrieve/pii/S0001691807001278.

Varazzani C, San-Galli A, Gilardeau S, Bouret S (2015) Noradrenaline and dopamine neurons in the reward/effort trade-off: a direct electrophysiological comparison in behaving monkeys. J Neurosci 35:7866–7877 Available at: http://www.ncbi.nlm.nih.gov/pubmed/25995472.

Ventura-Bort C, Wirkner J, Genheimer H, Wendt J, Hamm AO, Weymar M (2018) Effects of Transcutaneous Vagus Nerve Stimulation (tVNS) on the P300 and Alpha-Amylase Level: A Pilot Study. Front Hum Neurosci 12:202 Available at: http://www.ncbi.nlm.nih.gov/pubmed/29977196.

Vonck K, Raedt R, Naulaerts J, De Vogelaere F, Thiery E, Van Roost D, Aldenkamp B, Miatton M, Boon P (2014) Vagus nerve stimulation…25 years later! What do we know about the effects on cognition? Neurosci Biobehav Rev 45:63–71 Available at: http://dx.doi.org/10.1016/j.neubiorev.2014.05.005.

Warren CM, Tona KD, Ouwerkerk L, van Paridon J, Poletiek F, van Steenbergen H, Bosch JA, Nieuwenhuis S (2018) The neuromodulatory and hormonal effects of transcutaneous vagus nerve stimulation as evidenced by salivary alpha amylase, salivary cortisol, pupil diameter, and the P3 event-related potential. Brain Stimul 12:1–8 Available at: https://doi.org/10.1016/j.brs.2018.12.224.

Warren CM, Wilson RC, van der Wee NJ, Giltay EJ, van Noorden MS, Cohen JD, Nieuwenhuis S (2017) The effect of atomoxetine on random and directed exploration in humans. PLoS One 12:e0176034 Available at: http://www.ncbi.nlm.nih.gov/pubmed/28445519.

Waschke L, Tune S, Obleser J (2019) Local cortical desynchronization and pupil-linked arousal differentially shape brain states for optimal sensory performance. Elife 8 Available at: http://www.ncbi.nlm.nih.gov/pubmed/31820732.

Wilcoxon F (1945) Individual Comparisons by Ranking Methods. Biometrics Bull 1:80 Available at: https://www.jstor.org/stable/10.2307/3001968?origin=crossref.

Yakunina N, Kim SS, Nam E-C (2017) Optimization of Transcutaneous Vagus Nerve Stimulation Using Functional MRI. Neuromodulation 20:290–300 Available at: http://www.ncbi.nlm.nih.gov/pubmed/27898202.

Yanagawa T, Chao ZC, Hasegawa N, Fujii N (2013) Large-scale information flow in conscious and unconscious states: An ECoG study in monkeys. PLoS One 8:1–13.

Yap JYY, Keatch C, Lambert E, Woods W, Stoddart PR, Kameneva T (2020) Critical Review of Transcutaneous Vagus Nerve Stimulation: Challenges for Translation to Clinical Practice. Front Neurosci 14:284 Available at: http://www.ncbi.nlm.nih.gov/pubmed/32410932.

